# Evaluation of RNAi and CRISPR technologies by large-scale gene expression profiling in the Connectivity Map

**DOI:** 10.1101/147504

**Authors:** Ian Smith, Peyton G Greenside, David Wadden, Itay Tirosh, Ted Natoli, Rajiv Narayan, David E Root, Todd R Golub, Aravind Subramanian, John G Doench

## Abstract

The application of RNA interference (RNAi) to mammalian cells has provided the means to perform phenotypic screens to determine the functions of genes. Although RNAi has revolutionized loss of function genetic experiments, it has been difficult to systematically assess the prevalence and consequences of off-target effects. The Connectivity Map (CMAP) represents an unprecedented resource to study the gene expression consequences of expressing short hairpin RNAs (shRNAs). Analysis of signatures for over 13,000 shRNAs applied in 9 cell lines revealed that miRNA-like off-target effects of RNAi are far stronger and more pervasive than generally appreciated. We show that mitigating off-target effects is feasible in these datasets via computational methodologies to produce a Consensus Gene Signature (CGS). In addition, we compared RNAi technology to clustered regularly interspaced short palindromic repeat (CRISPR)-based knockout by analysis of 373 sgRNAs in 6 cells lines, and show that the on-target efficacies are comparable, but CRISPR technology is far less susceptible to systematic off-target effects. These results will help guide the proper use and analysis of loss-of-function reagents for the determination of gene function.

## Introduction

The Connectivity Map (CMAP), a component of the Library of Integrated Cellular Signatures (LINCS), aims to build a comprehensive look-up table of the mRNA expression consequences of perturbing a cell. Conceptually, the pattern of mRNA changes serves as a signature of the perturbation, and correlations between these signatures allows insight into connections between genes, drugs, and disease states [1-5]. The original CMAP dataset (build 01) examined 164 small molecules, and build 02 increased that number to over 6,000. Even with this limited-scale dataset, query of CMAP has provided biological insight across diverse disease areas [6-9]. Genetic perturbations are an attractive companion to small molecules, allowing the direct interrogation of gene function in order to understand how gene dysfunction leads to disease states. Here we provide the first systematic exploration of the use of both RNAi and CRISPR loss-of-function technologies in the context of CMAP.

Soon after its discovery as an endogenous biological process, RNAi became the leading technology for the disruption of any gene of interest, especially in mammalian systems that were previously refractory to genetic manipulation [10]. The combination of scalable reagent creation, facile cellular delivery, and potent gene knockdown has enabled genome-wide RNAi screens, leading to a wealth of information on gene function [11-13]. More recently, components derived from CRISPR loci in prokaryotes have been developed as an orthogonal means of perturbing gene function in mammalian cells [14-16]. In this system, the Cas9 nuclease is programmed with a single guide RNA (sgRNA) to create a targeted dsDNA break [17]. When this break occurs in protein coding regions, indels arising from the error-prone non-homologous end-joining pathway (NHEJ) can result in frameshift mutations and a null allele. Like RNAi, CRISPR technology can be used at genome-scale for phenotypic screens [18-21].

The specificities of RNAi and CRISPR technologies are a critical concern for the reliable interpretation of experimental results. That molecular triggers of the RNAi pathway, while designed to silence a single gene of interest, can in fact affect multiple off-target genes has long been documented [22-24]. Indeed, in some cases, independent RNAi screens ostensibly examining the same phenotype have reported widely different hit lists [25]. And while some of this difference may be due to variations in experimental conditions, it likely also reflects, in part, the contamination of hit lists by genes erroneously nominated due to off-target effects. In mammalian cells, where the trigger is usually a synthetic siRNA or a transcribed shRNA, there are two major modes of off-target activity. First, the RNA sequence intended to be the passenger strand can instead be selected by AGO2 as the targeting strand; in this case, all activity from this unintended loading of AGO2 will be off-target [26]. Fortunately, the rules determining strand selection have been well-studied and proper design can minimize this mode of off-target activity [27,28]. The second major mode of off-target activity, however, is not easily dealt with during the design of RNAi triggers: the small RNA intended to target a unique transcript can instead enter the miRNA pathway and potentially contribute to the silencing of dozens if not hundreds of transcripts [22,23]. Indeed, for endogenous miRNAs, both computational algorithms to predict mRNA targets and experimental approaches to detect them *en masse* have shown that many transcripts are subject to miRNA-based regulation [29,30]. While the mRNA targets of a miRNA are still not fully predictable, it is clear that the miRNA seed sequence – nucleotides 2 through 7 or 8, counting from the 5’ end – is a major determinant of activity.

Studies of CRISPR technology have shown that sgRNAs intended to target one specific genetic locus can lead to detectable cleavage of off-target DNA sites, although the reported promiscuity of Cas9 from *Streptococcus pyogenes* has varied widely [31,32]. Importantly, isolation of clones arising from CRISPR-mediated gene editing followed by whole-genome sequencing has shown that off-target rates can be reduced below the limit of detection, although this gold-standard assay was performed with only a small number of sgRNAs and is not practical for routine testing [33]. Nevertheless, these results offer the exciting possibility that, as understanding of the on- and off-target parameters of sgRNA design increases, CRISPR technology may be deployed with negligible off-target effects [34,35].

To build the CMAP dataset, 978 transcripts designated as landmarks are detected using Luminex bead-based technology, allowing high-throughput, low-cost data collection across many perturbations and cell types [36]. Here, we have analyzed the gene expression consequences of ~13,000 shRNAs profiled across 9 cell lines, allowing for a rigorous and deep exploration of the biological effects of RNAi reagents. Additionally, we generated gene expression signatures of CRISPR-Cas9 knockout with 373 sgRNAs in 6 cell lines, allowing a direct comparison of the relative potency and specificity of these loss-of-function technologies. Finally, we compared the gene-level results obtained for CRISPR and RNAi to each other. These results will guide the proper use and analysis of genetic perturbations and demonstrate the value of systematic, large-scale look-up tables such as the Connectivity Map.

## Results

### Widespread Off-target Effects with RNAi

RNAi reagents that do not have a sequence-matched target, such as scrambled sequences or those designed against reporter genes, are commonly used as experimental controls. By definition, on-target effects of a perturbation are the reproducible gene expression changes relating to the inhibition of the intended target gene, while off-target effects are all other reproducible changes specific to a reagent. From the CMAP dataset, we examined the correlations among signatures of 48 shRNAs targeting non-expressed control genes and observed that biological replicates of the same shRNA correlate while shRNAs of different sequences do not correlate (Fig. 1a, Supp. Fig. 1a, 1b). This correlation indicates reproducible gene expression changes that are specific to the shRNA perturbation despite the absence of any on-target activity, which implies the presence of a sequence-specific off-target signal.

**Figure 1.**
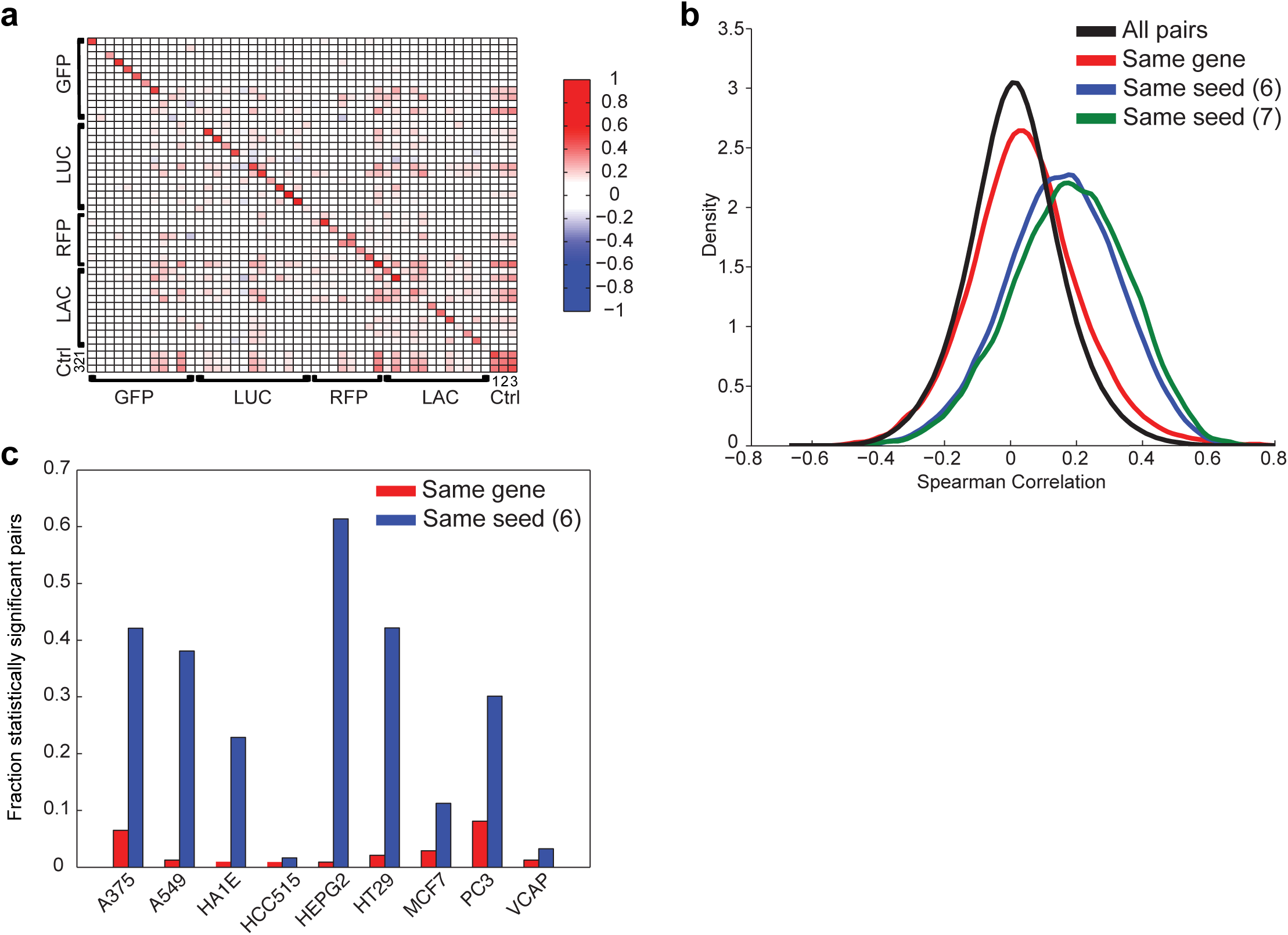
RNAi reagents have widespread off-target effects. a) Heatmap of Spearman correlations among pairs of shRNAs targeting control genes. Correlation on the diagonal reveals gene expression signal that is reproducible and specific to each shRNA despite the absence of a target. Control genes are labeled: GFP (green fluorescent protein); LUC (firefly luciferase); RFP (red fluorescence protein); LAC (beta-galactosidase). Additional control treatments are grouped under Ctrl; 1: pgw, a lentivirus with no U6 promoter and no shRNA; 2: empty_vector, a lentivirus with a run of 5 thymidines immediately after the U6 promoter, to terminate transcription; 3: UnTrt, wells that did not receive any lentivirus. b) Distribution of pairwise correlations of shRNA signatures with the same gene target (red), the same 6- and 7-mer seed sequence (blue, green), and all pairs of shRNAs (black). Data shown are from HT29 cells. Pairs of shRNAs with the same seed correlate much better than those with the same gene, which correlate only marginally better than random pairs. c) The fraction of pairs of shRNA signatures with the same target gene (red) or the same 6-mer seed (blue) that are statistically significant (q < 0.25) in each cell line. In all cell lines, correlation due to seed is more often significant than correlation due to gene.

Molecular triggers of the RNAi pathway are known to enter the miRNA pathway, where a seed sequence of 6 - 7 nucleotides (nts) directs the repression of off-target transcripts [22,23]. If shRNAs were entering the miRNA pathway to a significant degree, then we would expect the gene expression profiles of shRNAs sharing a seed sequence to correlate to each other. We examined all 7 nt windows along the shRNA and saw that shRNAs sharing a heptamer starting at positions 11 or 12 of the sense strand correlated strongly (Supp. Fig. 1c). These positions correspond to nts 2 – 8 of the antisense strand of the most-abundant Dicer-processing products of this shRNA design (Supp. Fig. 1d) [37], suggesting that most shRNAs do indeed enter the miRNA pathway and produce reproducible gene expression consequences, as has been observed previously [22,23,38].

We next compared the consistency of gene expression changes elicited by two classifications of shRNAs: i) shRNAs with different seed sequence that target the same gene; ii) shRNAs designed to target different genes but sharing the same seed sequence. In this set of 18,263 shRNAs, fewer than 1% share both a target gene and a seed sequence. Compared to the null distribution of all pairwise correlations, shRNAs targeting the same gene are more correlated to each other, but the increase in correlation is small compared to that of shRNAs sharing the same seed sequence (Fig. 1b). Interestingly, these results were more pronounced in some cell lines, although in all cases the correlation between same-seed pairs was greater than same-gene pairs (Fig. 1c). This finding means that a larger component of gene expression changes due to shRNA treatment are a consequence of the seed sequence rather than the target gene knockdown. This further suggests that the reliance on an individual shRNA for assessing the phenotypic consequences of gene silencing will often lead to erroneous conclusions, as seed sequence effects are prevalent.

### Consensus Gene Signatures

To attempt to accurately measure the on-target signature, we sought to combine gene expression information from multiple shRNAs targeting the same gene - but with different seeds - to produce a Consensus Gene Signature (CGS). As a first approach, we used a weighted average of multiple perturbations with weights based on a pairwise correlation matrix (Fig. 2a). To evaluate the fidelity of a CGS, we introduce the holdout method: divide at least six shRNA signatures targeting the same gene into two disjoint groups, make a CGS from each, and find the correlation of the two CGSs (Supp. Fig 2a). For genes with enough signatures to generate two disjoint consensus signatures (>=6), holdout analysis checks whether the two solutions agree better than nulls generated by permutation with cross-gene shRNAs. If on-target activity is a sufficiently large component of each CGS, then they will correlate with each other better than with a null population of CGSs made from random draws of shRNAs.

**Figure 2.**
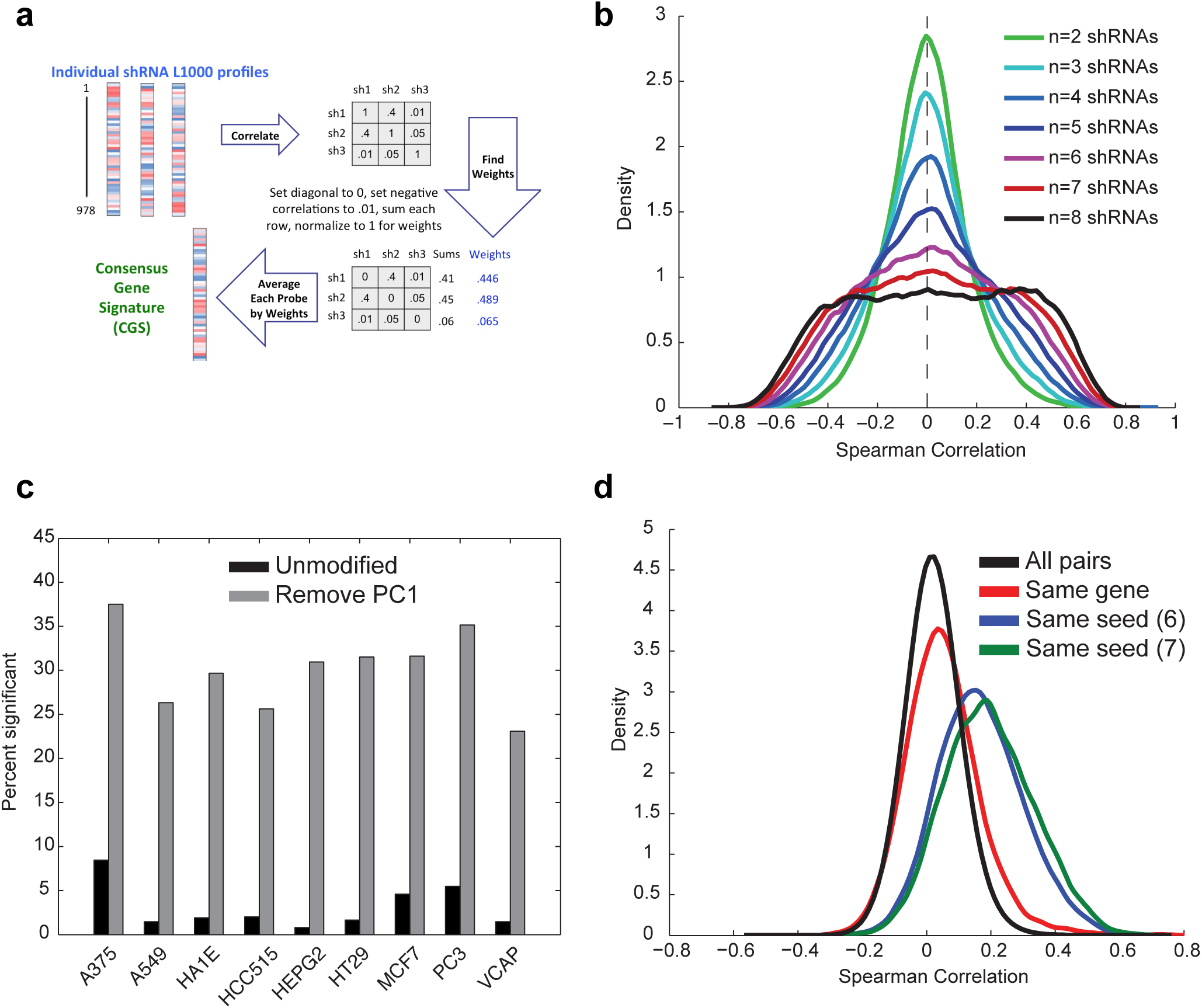
Removal of First Principal Component (PC1) improves assay signal. a) Schematic of the weighted average procedure for combining individual shRNA signatures targeting the same gene into a Consensus Gene Signature (CGS). The shRNAs are weighted by the sum of their correlations to other same-gene shRNAs and then averaged. b) CGSs made from random groups of shRNAs show increas-ing variance of Spearman correlation with larger numbers of component shRNAs. Because these are random groups, there should not be a consistent signal; the increasing probability of very large correlations reveals a spurious signal that we attribute to the first principal component of the data. c) The percentage of genes with 6 or more shRNAs that have statistically significant holdout results for each cell line with PC1 retained (unmodified, black) and PC1 removed (gray). d) Removal of PC1 does not diminish the magnitude of the seed effect. After removal of PC1, distribution of pairwise Spearman correlations in HT29 for pairs of shRNAs with same gene target (red), the same 6- and 7-mer seed sequence (blue, green), and all pairs of shRNAs (black).

Interestingly, examination of the permutation null CGSs revealed that increasing the number of component shRNAs led to a substantial increase in the mean and variance of correlation, that is, a generic increase in correlation implying a common, gene-target independent effect (Fig. 2b). Principal component analysis revealed that the first principal component (PC1) of the data was large, accounting for 10-20% of the total variance across both cell lines and perturbation types (i.e. for both shRNAs and small molecules in the CMAP dataset). Further, the fraction of variance accounted for by PC1 in CGSs increased as the number of input shRNA signatures increased in the perturbation nulls, suggesting that PC1 drives the spurious increase in correlation between null CGSs (Supp. Fig. 2b).

Supervised and unsupervised analysis of principal components has been previously used in gene expression analysis, surrogate variable analysis, and population genetics to characterize and remove unwanted variation [39,40], and thus we sought to test the effect of removing PC1 in these data. We performed holdout analysis and compared the results from keeping or removing PC1. Of 2,187 gene-cell line pairs with six or more shRNA signatures, only 2.9% were statistically significant (*q* < 0.25) with PC1 included, while 28.8% were significant with PC1 removed, and this dramatic improvement was observed across all cell lines (Fig 2c, Supp. Fig 2c). Notably, removal of PC1 did not lead to a decrease in the seed-effect, which is still much larger than the gene-effect (Fig 2d, compare to Fig 1b). Holdout analysis further suggests that cell context is pertinent for observing gene function, as most genes have a statistically significant holdout result in a fraction of cell lines, not all of them (Supp. Fig 2d). We speculate that PC1 may be a large, generic biological effect – e.g. an effect on cell cycle or viability – that obscures true biological connections, and that this effect is highly variable even among biological replicates of the same perturbation. Removal of PC1 followed by holdout analysis can thus be used to identify CGSs that are sufficiently enriched for on-target signal to be representative of the intended effect of gene transcript depletion.

To evaluate the effectiveness of the CGS for extracting on-target effects, we first examined the suppression of the target genes when they were directly measured in the expression signatures, i.e. targeting of landmark transcripts. We saw that 60.6% of shRNA target genes rank in the top 1% of most-downregulated transcripts when using the CGS, compared to 36.0% when using data from individual shRNAs (*p* < 10^-10^ by Kolmogorov-Smirnov test, Fig. 3a). When we removed PC1, the result did not change appreciably: 61.6% and 36.4% of target transcripts ranked in the top 1% with the CGS and individual shRNA, respectively. It is unsurprising that PC1 does not substantially affect the rank of the target knockdown gene because it is a broad signal with changes across many measured genes, whereas the knockdown of the target gene is large relative to that gene’s background distribution. Thus, in the special case where we know that transcript modulation is on-target, the CGS improves signal. We next evaluated the effectiveness of the CGS for the entire signature: using a leave-one-out approach, we found that 94% of individual shRNAs correlate better to the CGS made from remaining shRNAs than they do to other individual shRNAs targeting the same gene (Fig. 3b). By comparison, 77% of shRNAs correlate more strongly with an individual shRNA sharing the same seed than to the CGS containing that shRNA, indicating that use of the CGS partially mitigates the seed-based off-target signal of shRNA signatures (Fig. 3b).

**Figure 3.**
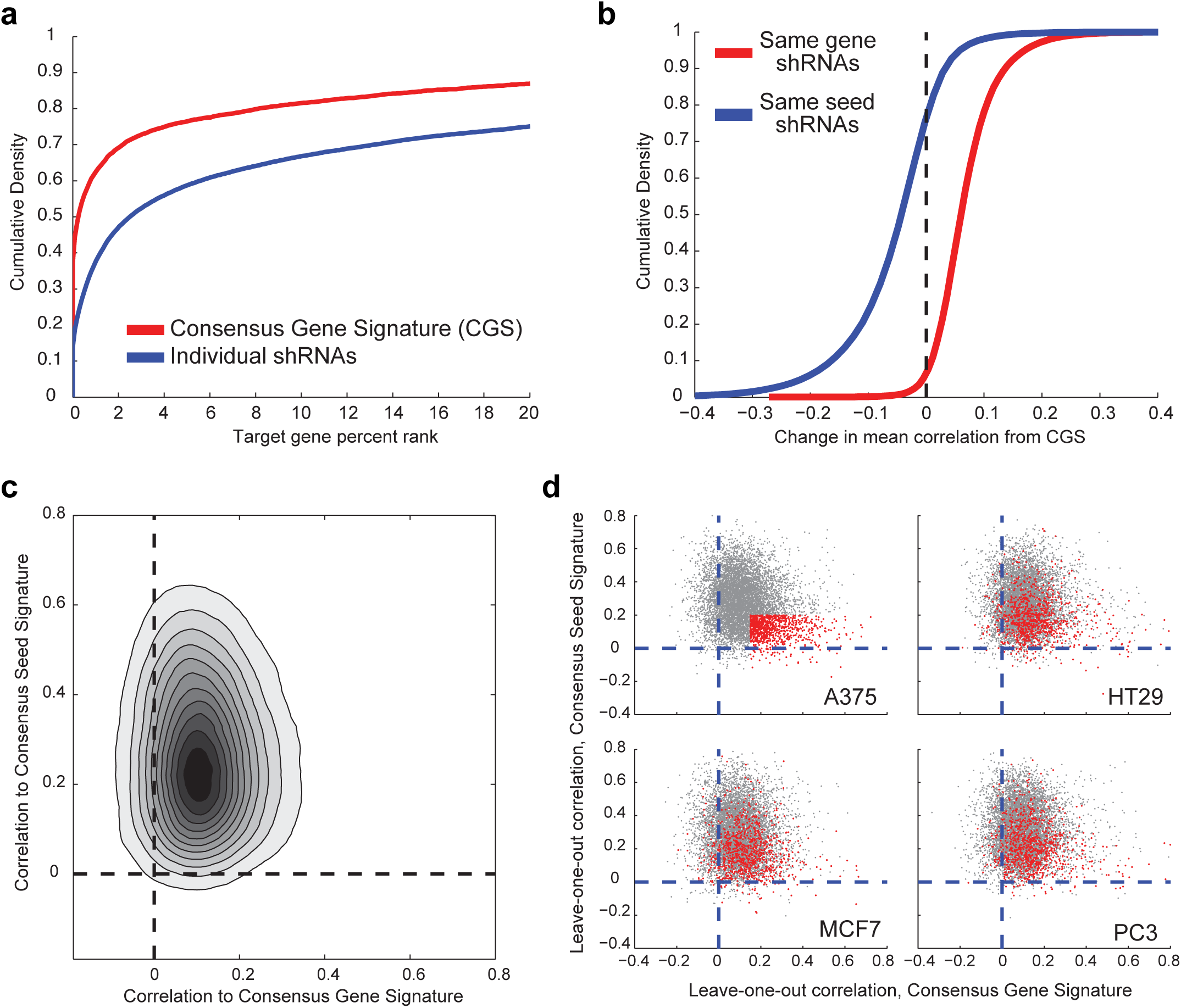
Consensus Gene Signature (CGS) enhances on-target signal and mitigates off-target effects. a) For directly-measured landmark transcripts targeted by RNAi, cumulative distribution of the rank of that transcript in the resulting signature when using either the signature produced by an individual shRNA (blue) or the CGS (red). b) Cumulative distribution plot of the change in the correlation to the CGS for either individual shRNAs that target the gene (red) or for shRNAs that share the same seed as one of the shRNAs that contributed to the CGS (blue). c) For individual leave-one-out shRNAs, comparison of the correlation to the CGS and the analogous Consensus Seed Signature (CSS). The density color scale is linear. d) Comparison of on- and off-target activity across cell lines. Top *left*: for each shRNA in A375 cells, plot shows the correlation between CGS (x-axis) and CSS (y-axis). Those in red have minimal off-target effects (CSS Spearman correlation < |0.2|) and substantial on-target effects (CGS Spearman correlation > 0.15). *Remaining panels*: The red-highlight shRNAs from A375 cells are highlighted in red in three other cell lines.

To further examine individual shRNAs, we used the leave-one-out approach to create a CGS from all remaining shRNAs targeting the same gene and an analogous Consensus Seed Signature (CSS) from shRNAs with the same seed sequence but different gene targets. We observed that the majority of shRNAs exhibit both on- and off-target activity – quantified by the Spearman correlation of the shRNA signature to the CGS and CSS, respectively (Fig. 3c). Interestingly, in any particular cell line, a minority of shRNAs exhibited substantial on-target activity with minimal off-target activity. While the activities of these shRNAs showed variability in other cell lines, the performance across cell lines was correlated (Fig. 3d, mean odds ratio of the pairwise contingency tables of 5.6 and *p* < 10^-10^ by Fisher’s exact test). Thus, while shRNAs that are predominantly on-target in one cell line are more likely to be predominantly on-target in another, individual shRNAs that provide an on-target phenotype in one cell context must be re-confirmed in other cellular contexts. The use of explicit seed-matched controls has proven useful to experimentally assess the seed contribution to a phenotype [41].

Finally, we asked whether the magnitude of gene expression changes due to a particular seed sequence was consistent across cell lines. For each seed sequence, we calculated the effect magnitude by computing the mean leave-one-out CSS correlation across all shRNAs with that seed for each cell line. We then compared the set of cell line means for each seed to the collection of all seed cell line means, with the null hypothesis that each seed has the same distribution of mean effect magnitude across cell lines. Using the F-test, we found that 66% of seeds had smaller variances of mean leave-one-out correlation than the bulk population (*q* < 0.25), implying that some seed sequences have similar magnitudes of effect across multiple cell lines (Supp. Fig. 3a) – i.e. that some seeds induce consistently small or large gene expression changes across lines. By comparison, 16% of genes had smaller variance (q < 0.25) than the bulk population (Supp. Fig 3b). This analysis reveals that many seeds show consistent magnitude of effect across cell lines, and that it may be possible to predict whether a particular shRNA will be predominantly off-target using empirical data from other cell lines. This also suggests that certain seeds – those with consistently large gene expression changes – should be avoided in shRNA library design, while those seeds with smaller effect sizes should be preferred.

### Evaluation of CRISPR-Cas9 signatures

CRISPR technology provides an alternative means of examining the effect of gene silencing. To measure the performance of sgRNAs in gene expression space, we used on-target activity rules [42] to design 6 or 7 sgRNAs against 53 genes, 20 of which were directly measured landmark transcripts, along with negative controls targeting EGFP, beta-galactosidase, and luciferase, for a total of 373 sgRNAs. These were profiled at 96 hours post-infection in six cell lines engineered to express Cas9. To directly compare RNAi results to the CRISPR dataset, we also evaluated from the CMAP database the 395 unique shRNAs targeting the same genes in the same cell lines, for a total of 3,568 matched signatures. 50 of the 53 target genes from the CRISPR dataset had analogous RNAi data measured at the same time point and the same cell line; we note that this set of genes was biased towards those that showed evidence of a gene expression signature when assessed by RNAi.

We first investigated the change in mRNA of the targeted landmark genes perturbed by RNAi and CRISPR technologies by measuring the z-scores for the differential expression of the target gene relative to the population (Fig. 4a). We observe comparable reduction in the transcript abundance of the target gene from the two technologies. Presumably, while mRNA is not the direct target of an sgRNA, the nonsense-mediated decay pathway is reducing the steady-state level of mRNAs carrying frameshifts caused by Cas9-mediated cleavage and NHEJ-mediated repair. We next employed holdout analysis to gauge the efficacy and consistency of sgRNAs, as previously done with shRNAs, including a comparison of results with and without PC1. The first principal component direction among sgRNA signatures was essentially the same as among shRNAs – the two have a Spearman correlation of 0.829 in the landmark basis. For the CRISPR dataset of 324 combinations of genes and cell lines, among the unmodified signatures, 33.6% were found to have significant correlation by holdout (*q* < 0.25), while 58.3% of holdout results were significant after removal of PC1, further showing that the performance improvement gained by removal of PC1 is not restricted to shRNAs (Fig. 4b). In the gene-matched shRNA data, of 297 gene-cell line pairs, 183 had 6 or more independent signatures, which is the minimum necessary to run the holdout analysis; of these, 24.6% and 47.0% were found to have statistically significant correlation with PC1 kept or removed, respectively (Fig. 4c). These results show that both RNAi and CRISPR technologies can generate reproducible, on-target results.

**Figure 4.**
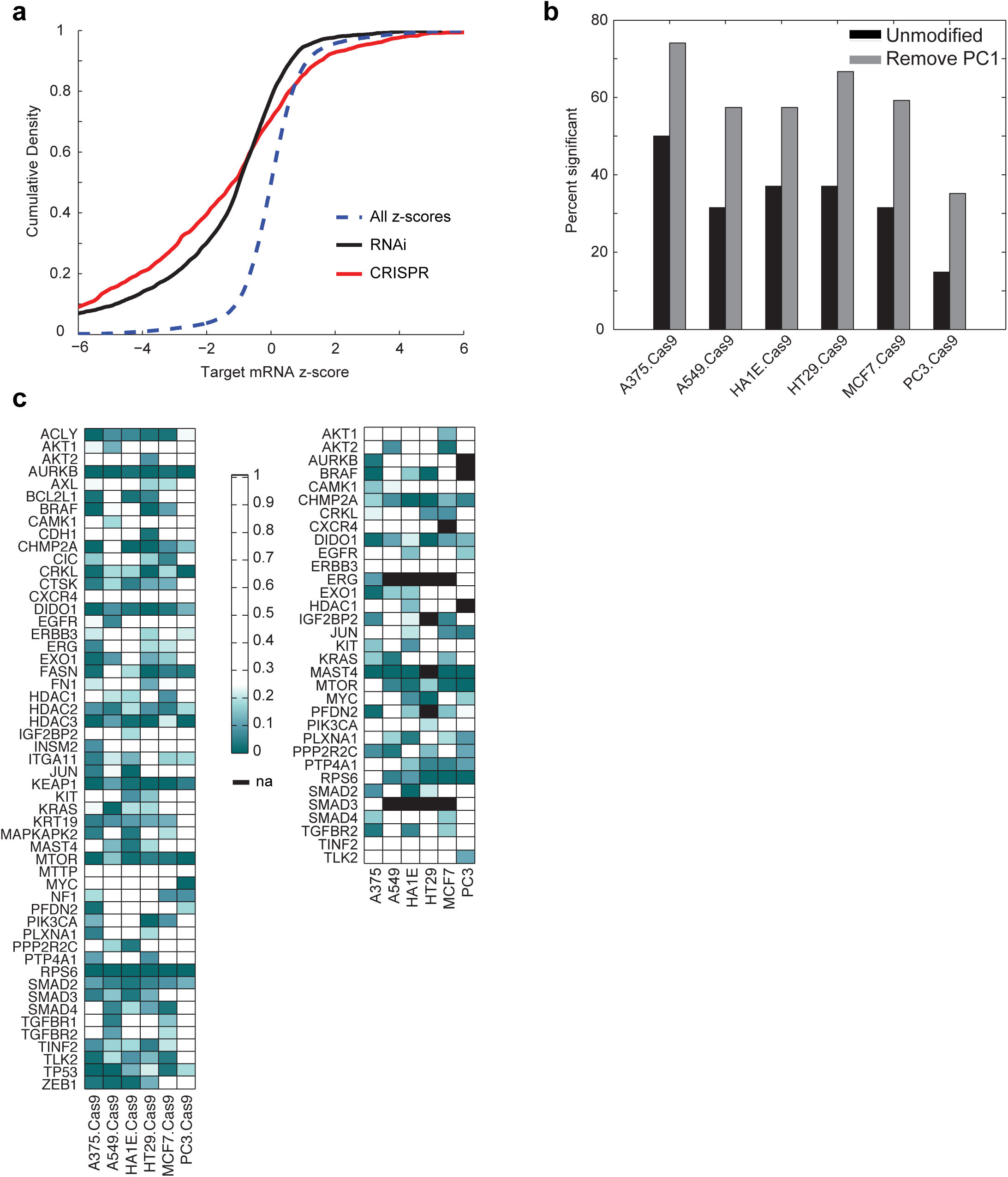
Gene expression analysis of CRISPR-Cas9 reagents. a) Analysis of landmark transcript reduction, comparing CRISPR and RNAi for genes targeted by both technologies. The dotted line (blue) is a null distribution of the set of all z-scores. Both technologies show significant downregulation of directly measured target transcripts. b) In the CRISPR data, comparison of the fraction of genes with significant holdout results in each cell line with or without removal of PC1 (q < 0.25). Removing PC1 from the dataset improves the number of statistically significant genes across all cell lines. c) Holdout analysis for genes assayed by CRISPR (left) and RNAi (right). Because holdout analysis requires at least six independent reagents, not all of the genes assayed by CRISPR have sufficient coverage by RNAi; missing values are indicated by black boxes. Smaller q-values (green) indicate greater statistical significance, i.e. that the Consensus Gene Signature is valid.

A large number of DNA cuts by CRISPR-Cas9 in a cell can produce viability effects, either because the target site is amplified or because of cutting at off-target sites [43,44]. We investigated systematic gene expression changes due to copy number amplification by training a linear model to predict copy number of target genes. We first attempted to use the hallmark DNA repair gene set from MSigDB, but observed no enrichment of that gene set with increasing copy number [45]. Because the measured landmark gene set contains only a fraction of genes, it may be too small to show gene set enrichment. To search for a copy number signal with an unbiased approach, we trained a lasso-regularized linear regression model using the set of all reproducible sgRNA signatures to predict copy number annotations from the Cancer Cell Line Encyclopedia [46]. Using 20-fold cross validation, we found a small but statistically significant correlation; the predicted vector had a Pearson correlation of 0.30 with the copy number data (*p* < 4x10^-15^).

Adding credence to the model, non-targeting control sgRNAs had less variance along this axis than did reproducible sgRNAs with a gene target (*p* = 6x10^-6^ by the f-test). Importantly, the variance explained by this axis is about ten-fold smaller than the variance explained by on-target activity of reproducible sgRNAs; in the future, application of a non-linear model to a larger data set may be informative to better quantitate this effect. This result suggests that there is indeed a copy number signal in gene expression space that should be considered when interpreting experimental results but that, in most cases, it is small enough that it should not typically obscure on-target gene signal.

### Projection Analysis

Unlike the seed-based off-target effects of shRNA reagents, which comprise a large fraction of the gene expression change, the penetrance of systematic off-target effects of sgRNAs is unknown. To interrogate the magnitude of on- and off- target effects in both sgRNA and shRNA signatures, we formulated a novel decomposition analysis using projection. A gene expression signature is a combination of three components: on-target activity, off-target effects, and assay noise. Projection estimates on-target activity by comparing the signature of an individual perturbation to signatures from other perturbations targeting the same gene, and quantifies off-target activity by measuring the similarity among replicates of a signature after subtracting the on-target activity. Because the method estimates off-targets by searching for reproducible elements of the signature which are not related to on-target activity, it does not require prior knowledge of the off-target mechanism to accurately detect off-target signal.

We first assessed the consistency of projection decomposition to detect on-target effects. Of 2,231 sgRNA signatures, 1,723 individual sgRNAs (77%) had significant on-target activity (*q* < 0.25) by this analysis; for the matched 2,183 shRNA signatures, 931 (43%) were significant. To evaluate whether projection corroborates holdout analysis - an alternate method for measuring on-target activity – we then examined signatures belonging to genes with statistically significant holdout results, and saw these were preferentially enriched by projection: 91% of individual sgRNAs and 61% of individual shRNAs from genes with significant holdout results were found to have significant on-target projection ranks. Because the holdout method requires two disjoint sets to arrive at the same answer, it is stringent, requiring a majority of the signatures to demonstrate on-target activity. Therefore, individual signatures with on-target activity are necessary but not sufficient to give a statistically significant holdout result.

We next examined the ability of projection decomposition to reveal known off-target effects, in this case, the miRNA-seed effect in shRNAs. For each shRNA, we compared its projection estimate of off-target activity to its correlation to the leave-one-out CSS, and found a significant relationship (Pearson = 0.4275, *p* < 10^-10^, Fig. 5a). While these two metrics are not strictly identical, their correspondence indicates that decomposition by projection - by modeling on-target activity with independent perturbations and off-target effects with assessment of technical replicates - can reveal off-target effects without prior knowledge of their source.

**Figure 5.**
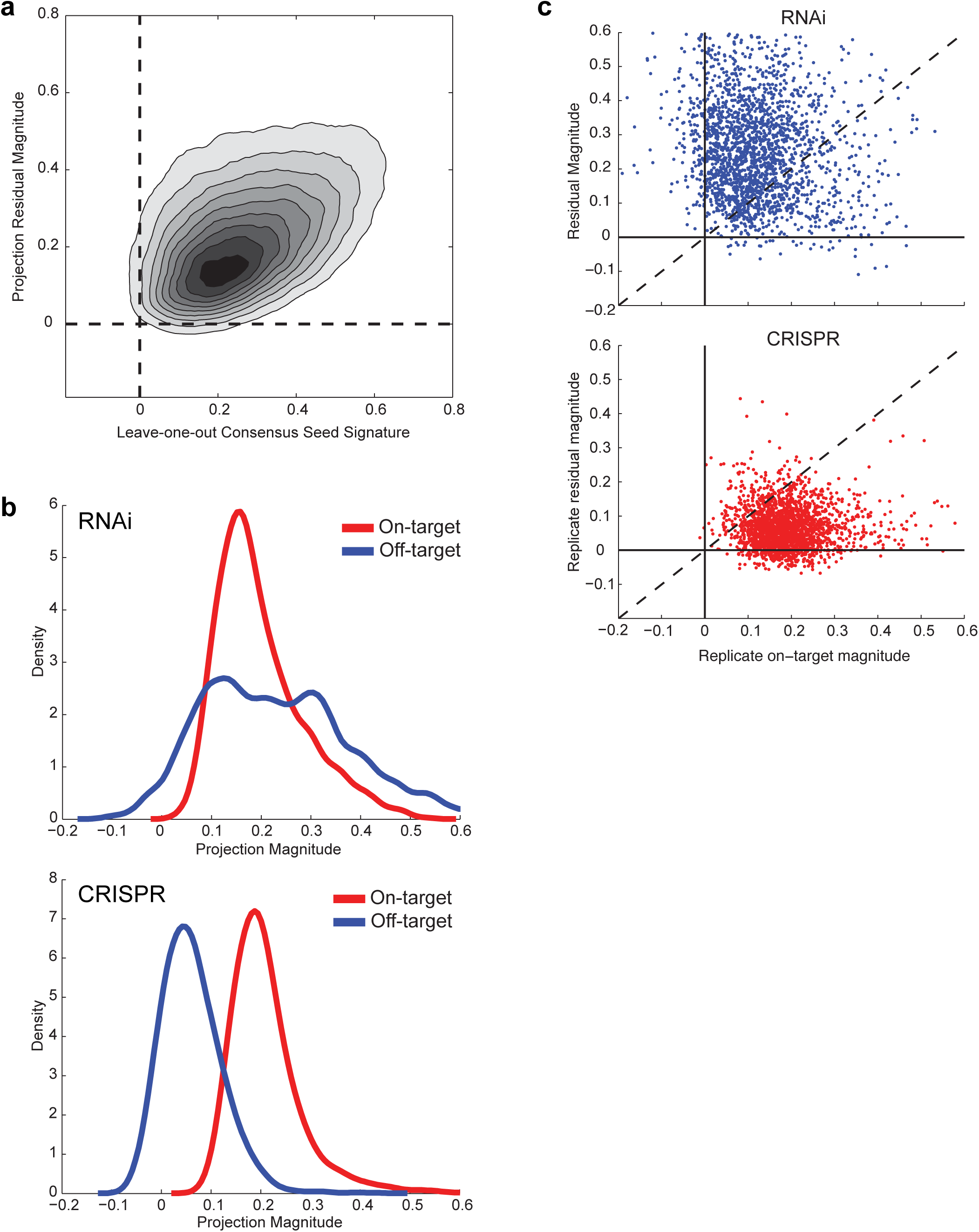
Decomposition by projection. a) For shRNAs, magnitude of off-target effect com-paring the leave-one-out Consensus Seed Signature to projection. Pearson correlation coefficient = 0.43. b) For individual shRNAs (top) and sgRNAs (bottom) where the on-tar-get magnitude passes FDR < 25%, distribution of on- and off-target magnitudes as assessed by projection decomposition. c) Scatter plots of on-target and off-target projection magnitudes for RNAi (top) and CRISPR (bottom) for all of the signatures of reagents in the data set. While the two technologies show similar on-target activities, RNAi shows large off-target effects.

We then compared the relative magnitude of on- and off-target effects for RNAi and CRISPR reagents. Considering only the 1,723 sgRNA signatures with statistically significant on-target activity, we observed an average on-target magnitude of 0.211 and off-target magnitude of 0.062, with 97.4% having a larger on-target than off-target component (Fig. 5b, c). For the 924 shRNAs targeting the same genes with significant on-target projection ranks, the mean on-target magnitude was comparable to sgRNA-based knockout, 0.197; however, the mean off-target magnitude was 0.230, and only 41.8% shRNAs had a larger on-target than off-target component (Fig. 5b, c). Thus, while RNAi reagents necessarily engage the miRNA pathway to cause off-target effects, these results suggest the exciting possibility that CRISPR technology can frequently produce faithful on-target signatures with low levels of off-target signal.

### Connections between RNAi and CRISPR

We compared the signatures produced by RNAi and CRISPR technology to each other utilizing query-based metrics, as this is, ultimately, the most important measure of the quality of these data in the Connectivity Map. For each gene in each cell line, we created a CGS using all sgRNAs and queried the CMAP database of ~4,000 shRNA-derived CGSs. Of 297 sgRNA CGSs for which there was a same-gene shRNA CGS, 116 (39%) had statistically significant connectivity (*q* < 0.25, Fig. 6a). Further, when we require that both the sgRNA and shRNA CGSs separately pass holdout analysis, 74% (37/50) connect with *q* < 0.25 (Fig 6b). These results show that for many of the genes examined here, knockdown and knockout produce similar signatures. Interestingly, we find examples where holdout analysis suggested the resulting CGSs are an accurate measure of the on-target effects of gene knockdown and gene knockout, but these fail to find each other in a query, such as SMAD2 in three different cell lines; these may be cases where there are phenotypic differences as a function of gene dosage. It is also possible that different kinetics of gene depletion between the two technologies may affect connectivity results, or that the signatures could be false positives from the holdout analysis. These results suggest that, going forward in the Connectivity Map, CRISPR technology should be the first choice for conducting loss-of-function studies, with RNAi used as a complementary or secondary assay. The high on-target activity and low prevalence of off-target effects will enable the use of smaller numbers of perturbations per gene with CRISPR technology, which in turn will allow greater coverage across both genes and cell types.

**Figure 6.**
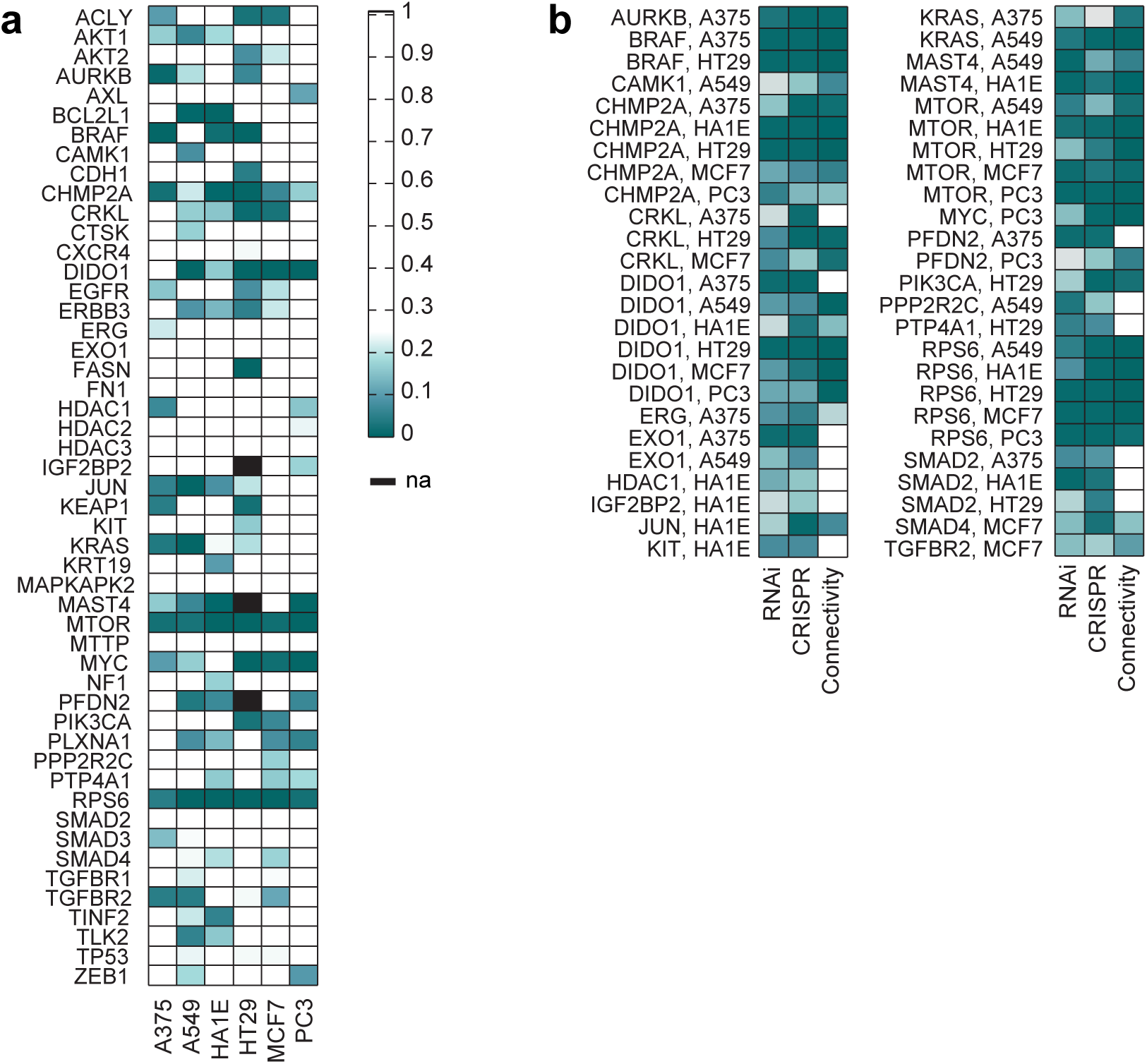
CMAP queries for RNAi and CRISPR reagents. a) For all genes assessed by both CRISPR and RNAi technologies, the q-values for connectivity across cell lines. b) For genes passing holdout analysis (q-values < 0.25) by both technologies individually in a cell line, the q-values for connectivity. Holdout analysis q-values are plotted for each technology in the first two columns, connectivity q-values are plotted in the third column.

## Discussion

For over a decade, RNAi has been widely used to induce gene suppression, especially in mammalian cells. More recently, CRISPR technology has emerged as an exciting new tool for loss-of-function perturbational studies. Using RNAi and CRISPR datasets generated as part of the Connectivity Map, we have characterized the gene expression changes induced by these two perturbation types, and shown that the on-target efficacy is similar. Analysis of off-target activity, however, shows that miRNA-like seed effects of RNAi are widespread, while CRISPR technology produces few systematic off-target effects. In many cases, we observe that RNAi and CRISPR produce comparable signatures to each other once non-specific and off-target effects are mitigated.

Analysis of large-scale gene expression data required the development of several analytical approaches. We have introduced the notion of a Consensus Gene Signature to combine information from multiple independent perturbations, as is commonly done in the analysis of genetic screens using these technologies: concordance of multiple perturbations of different sequence targeting the same gene suggests that the observed phenotype is due to on-target silencing. While the CGS approach is effective, future work will refine the CGS concept, as has been done with small molecule gene expression profiles [47], including using inference to model the on- and off- target components of a signature. Likewise, we developed holdout analysis to assess the reliability of a CGS. This approach, however, is stringent, requiring consistent on-target effects among a majority of the individual perturbations targeting the same gene; while disjoint CGSs might fail to correlate, the CGS from the entire set of perturbations could still be valid even if only a minority of perturbations are effective. Finally, we used projection to quantitate the magnitude of on- and off- target effects of a perturbation. This method is powerful because it does not require prior knowledge or characterization of off-target mechanisms to observe their effect. In the course of these analyses, we observed a large principal component in the dataset, irrespective of perturbation type and found that removal of PC1 led to improved results by several metrics. While unexpected in this dataset, removal of principal components to improve signal has precedent in many circumstances, for example GWAS and image processing. For large scale, high-throughput gene expression data sets, employing and improving these methods will be critical to discriminate valid gene perturbation signatures.

This is an exciting time for loss-of-function screens with genetic perturbations. RNAi experiments, when properly controlled and analyzed, have a proven record of generating discoveries. The prevalence of off-target effects detected at genome scale with RNAi, however, necessitates careful interpretation of results to accurately determine loss-of-function phenotypes - in particular, either computationally or experimentally accounting for the penetrance and heterogeneity of seed-based effects. Further, these results emphasize the long-standing recommendation on the use of multiple reagents to gain confidence in a reproducible phenotype. CRISPR technology, when analyzed by the same metrics for consistency of reagents targeting the same gene, tends to produce predominantly on-target activity. Potentially, agreement among a small number of independent sgRNAs may be sufficient evidence of on-target activity in a majority of cases - an important consideration when it comes to building a genome-wide map across many cellular contexts of the expression consequences of loss-of-function reagents. The proper use of these two technologies can synergistically bring new value to the use of genetic perturbations for gene function discovery and therapeutic development.

## Materials and Methods

### Data availability

The portal to access Connectivity Map data is https://clue.io. This website also provides detailed information on the experimental protocols used to generate these data. Additional information can be found at the Library of Network-Based Cellular Signatures (LINCS) Program website, http://www.lincsproject.org.

### L1000 Platform

In brief, the L1000 Luminex assay explicitly measures 978 specific mRNA transcripts designated as landmarks. Samples are run in 384 well plates, and the expression values are normalized z-scores across the distribution of expression values for each gene across a single plate. A signature of a perturbation – a small molecule, an shRNA, an sgRNA, or an ORF – is an average of the 978 differentially expressed transcripts of three biological replicates of cells treated with that perturbation. The signature consists of 978 z-scores indicating gene expression changes in the measured landmark genes due to treatment by the perturbation. Similarities between signatures were evaluated using Spearman correlation between z-scores in only the landmark space of 978 genes.

### Data Curation

We performed analyses on experiments from eight cancer cell lines (A375, A549, HCC515, HEPG2, HT29, MCF7, PC3, VCAP) and one immortalized cell line (HA1E); all lines were lysed 96 hours post viral infection except for VCAP, which was lysed at 120 hours to account for its slower growth rate. All shRNAs were run in triplicate, with each specific shRNA L1000 signature representing a weighted average between the three technical replicates.

### Relative On- and Off-Target Strengths

To quantify the relative strength of on-target and off-target effects, we compared the distribution of Spearman correlations for all pairs of shRNAs sharing the same target gene and all shRNAs sharing the same 6-mer or 7-mer seed sequence compared to correlations of all pairs of shRNAs. Within each cellular context, we identified all pairs of shRNAs that share either seed sequence or target gene and take a rank-based Spearman correlation between their expression profiles. We compared the distribution of same-seed shRNA correlations both for the 6-mer and 7-mer definitions of seed region and found comparable effects.

### False Discovery Rates (FDR)

To correct for multiple hypothesis testing, we use the Storey’s FDR correction to calculate a q-value given the p-values. First, we assign p-values in general by comparing observed statistics to permutation nulls. For example, in holdout analysis, we calculate the mean correlation between two consensus signatures iteratively generated from a set of signatures targeting the same gene. We generate a null distribution by applying the same procedure to permuted random groups, and we assign p-values to the measured values by ranking relative to the permutation null values. The interpretation of Storey’s q-value is that from a large collection of hypotheses, the set of hypotheses (numbering N) with q-value < t will have at most a fraction t of type 1 errors or false positives. That is, on average under the null hypothesis modeled by the permutation nulls, the expected number of type 1 errors is at most N*t. We use a q-value threshold of 0.25 for most of our analyses; as CMAP is designed to be a screening tool leading to hypotheses for follow-up experiments, we are willing to tolerate a relatively higher number of false positives.

### First Principal Component

The analysis of the principal components of the data used the standard PCA Matlab code, which in turn uses SVD to linearly transform the data into a new orthogonal basis. The data that was used in the analysis is the differentially expressed genes – the z-scored signatures – in which the means of each gene over all signatures are approximately zero, and the sign and magnitude in a signature indicates the response of that gene to the perturbation relative to the population. Note that PCA first mean centers the data, i.e. the mean of each gene is set to 0 before calculating the variance. In the analysis, we initially ran PCA on each cell line and perturbation type independently, though our final analysis considered the dataset as a whole. The first principal component (PC1) is the linear projection of the data with maximal variance – the axis with the greatest variation in the data. The weighting of PC1 for a signature is the magnitude of the signature in the first principal component direction, and by definition, the PC1 weighting across the signatures has larger variance than any other principal component weighting.

Our findings from PC1 are surprising for several reasons. First, the direction of the PC1 – its representation in landmark gene space – is virtually identical across cell lines and perturbation types. The direction of PC1 among small molecule treated A375 signatures is the same as that among shRNA signatures in MCF7, for instance. This suggests that independent of treatment and cell line, there is an axis along which genes are consistently modulated – implying a generic response. The weightings of PC1 are surprisingly large, accounting for 10-20% of the variance of signatures. Furthermore, the weights of PC1 increase among consensus signatures – in particular, the fraction of variance from PC1 increases even among null consensus signatures as more shRNAs are averaged together. This necessitates using a size-matched permutation null distribution – with the same number of shRNAs in each CGS - for each size of CGS, but it also suggests that the increase in correlation variance among CGSs of increasing size may be due to PC1.

Removing the PC1 is straightforward. First, we ran PCA on the entirety of the CMAP data – constituting over 400k signatures and use the PC1 from this global calculation as the axis on which all signatures were projected. The alternative would be to remove the local PC1, but while our analysis has shown this to be very consistent, running PCA on smaller data sets would introduce batch specific uncertainty in the PC1 removal. To remove PC1, the dataset is mean-centered, the component of the signatures in the direct of PC1 is subtracted, and the mean is added back on. The signal may be further improved by mean centering as part of the normalization process – as opposed to median centering – but this possibility has not been exhaustively explored. Finally, the direction of PC1 depends on both normalization and the calculation of differential expression. Our approach computes the z-score relative to each measured gene independently, so it should be equivalent to the quantile normalized data if each gene is scaled to variance 1.

### Consensus Gene Signature

To generate a consensus gene signature (CGS), we group all shRNA signatures targeting the same gene within the same cell line and apply the modz algorithm. We create a pairwise Spearman correlation matrix between the expression profiles of all signatures in this group, explicitly setting the diagonal of the correlation matrix to 0. The weight assigned to each shRNA signature is given by the sum across its corresponding row of the correlation matrix, with the weights normalized to sum to 1. The CGS is then given by a linear combination – or a weighted average – of the shRNAs, with coefficients given by the weights.

Target Gene Rank: To evaluate the rank of the target gene, we considered only the shRNA signatures where the targeted gene was measured explicitly as one of the 978 landmark genes. For each of these signatures, we ranked the 978 genes from most down-regulated to most up-regulated. We identified the rank (1 to 978) of the targeted gene and plotted the distribution of target gene ranks for each individual shRNA signature. We then formed consensus gene signatures from the individual shRNAs, and similarly identified the rank of the specifically targeted gene. We plotted the cumulative distribution of the individual shRNA target gene ranks compared to the consensus gene signature target gene ranks as a function of percent rank.

CGS-shRNA Correlations: We evaluated the enrichment of on-target effects by evaluating whether an individual shRNA signature correlates better to a disjoint CGS than to individual shRNAs used to create that CGS. For each gene and cell line with 2 or more shRNA signatures, for each signature in the group, we generated a CGS using all but one of those signatures. We compared the correlation between the CGS and the excluded shRNA to the mean correlation between the excluded shRNA and the shRNAs comprising the CGS. The difference between these two quantities is the improvement from using a CGS compared to a typical individual shRNA signature.

Holdout Analysis and CGS-CGS Correlations: We evaluated the enrichment in overall correlation between CGSs for the same gene. Within each cell line, we partitioned the shRNAs into 2 groups for genes with six more shRNAs. We generated two CGSs with the two groups and took a correlation between those two consensus signatures. We partitioned the groups 30 times and re-calculated the CGS correlation to avoid bias by a particular combination of shRNAs and summarized these partitions by taking the median correlation. To mitigate any artificial increases in-group correlation introduced by the CGS process, we introduced a permutation null by generating 10,000 identically sized groups of shRNAs targeting different genes and performing the same partition correlation calculation. For each gene, we assigned a p-value of enrichment by counting what fraction of permutation null groups had higher median correlation than the observed median correlation for that gene and cell line. To correct for multiple hypothesis testing, we assigned q-values using Storey’s FDR procedure and considered significant groups with q-values < 0.25. A gene in a cell line with a statistically significant holdout result indicates that independent CGSs agree better than would be expected assuming no common on-target effects among the shRNAs. Such a result indicates that the CGS of all of the shRNAs is representative of the on-target gene expression consequence of the target gene knockdown.

### Leave-One-Out Consensus Signature Correlations

We identified the group of shRNAs with at least one same-gene shRNAs and at least one same-6mer-seed shRNAs. For each of these shRNAs, we generated a consensus gene signature with the same-gene shRNAs, and a consensus seed signature with the same-6mer-seed shRNAs (excluding the shRNA itself in both groups). We found the Spearman correlation between the shRNA and its CGS and the shRNA and its CSS. We plotted the overall shRNA-CGS correlations on the x-axis and the shRNA-CSS correlations on the y-axis to estimate the magnitudes of the on- and off-target effects within a given shRNA.

### Projection Analysis

The projection method decomposes a signature of a perturbation into two orthogonal components representing the on-target and reproducible off-target effects without incorporating information about the mechanism of the off-targets. The most important outputs from projection are two quantities representing the normalized length of the on-target component and reproducible off-target component of the unit vector of the signature. First, a reference signature is constructed from other perturbations with the same purported on-target – i.e. a CGS of other shRNAs or sgRNAs targeting the same gene. We calculate the cosine similarity of the given signature to the reference and assign p-values by ranking that similarity relative to a number of permutation null reference signatures. The permutation nulls are CGSs constructed from unrelated groups of the same number of shRNAs or sgRNAs. For the projection magnitudes to be meaningful, the signature must have statistically significant similarity to the reference (*q* < 0.25). Without statistically significant similarity, either (1) the signature lacks measurable on-target activity or (2) the reference signature is not a good estimate of the true on-target signature. In the latter case, the true on-target component of the test signature will be incorrectly assessed as reproducible off-target.

If the test signature has significant similarity to the reference, residuals are calculated from the test signature’s replicates by projecting the replicates onto the subspace orthogonal to the reference signature. The on-target magnitude is given by the mean of the length of the replicates projected onto the reference divided by the length of the replicates; equivalently, the on-target magnitude is the mean inner product of the replicate unit vectors with the unit vector reference in the standard landmark basis. The off-target magnitude is given by the mean of the pairwise inner product of the replicate residuals normalized by the length of the replicates. It follows from the Pythagorean theorem that for *a* the on-target magnitude and *b* the off-target or residual magnitude, that *a*^2^ + *b*^2^ < 1. In fact, *a*^2^ + *b*^2^ for a given signature is identically the fraction of the signature that is reproducible, and this quantity correlates very well with the standard CMAP metric of reproducibility for a signature – the 75^th^ quantile of Spearman correlation among replicates of that signature (Supp. Fig 4).

The interpretation of projection is straightforward: any reproducible component of a signature orthogonal to the on-target activity is an unintended off-target – whether systematic biology, batch effect, or other artifact. The reproducibility of off-targets can be measured by considering the components of replicates of a signature orthogonal to an estimated reference. In general, we expect small off-targets because of batch effects and errors in calculating the reference signature – the on-target activity is not perfectly known, which is the original problem. That the calculated off-target activity correlates with the magnitude of the known seed effects in shRNA signatures as measured by correlation with the CSS validates the approach.

## Acknowledgements

We thank Oana Enache, Roger Hu, Larson Hogtrom, Jackie Rosains, Mudra Hegde, and David Lahr (Broad Institute) for analytical insight and helpful conversations. We thank the entire Genetic Perturbation Platform (GPP) and Connectivity Map (CMAP) teams for support and advice.

---

**Supplemental Figure 1.**
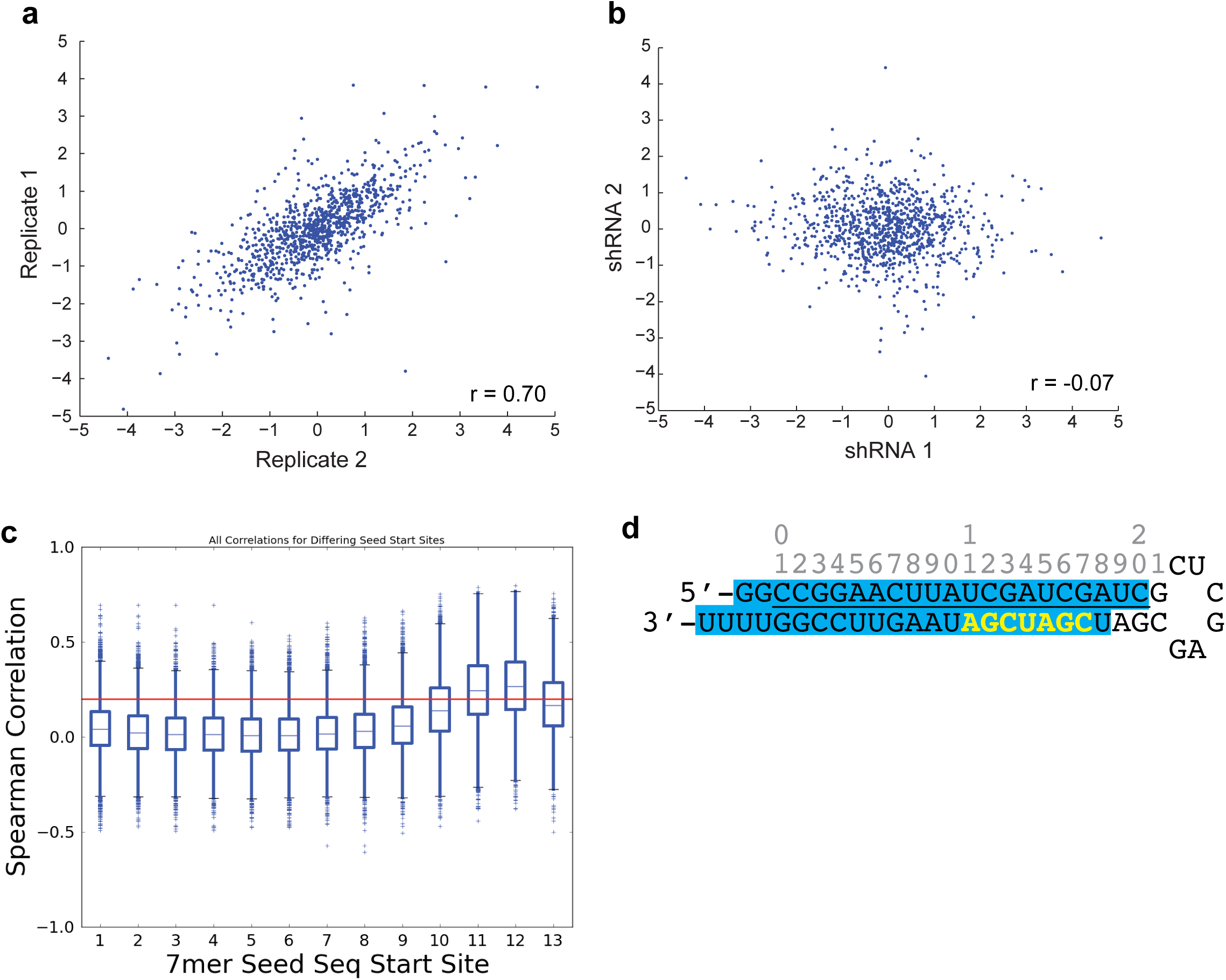
Control shRNAs and the seed effect. a) An example of strong correlation: two replicates of the same control shRNA show strong reproducibility as measured by the Spearman correlation of the differential expression of the 978 measured landmark genes. Each point represents one landmark transcript. b) An example of no correlation between signatures of different shRNAs. c) Schematic of shRNAs used in CMAP. The 21nt sense strand, which has the same sequence as the target mRNA, is underlined and numbered. The blue highlight indicates the major siRNA product produced after Dicer processing. The bolded, yellow nts indicate the seed sequence of the antisense / targeting strand. d) 7-mers beginning at positions 11 and 12 of the annotated sense strand show the greatest correlation, corresponding to the seed sequence of the antisense/targeting strand, and reflecting heterogeneity of Dicer processing.

**Supplemental Figure 2.**
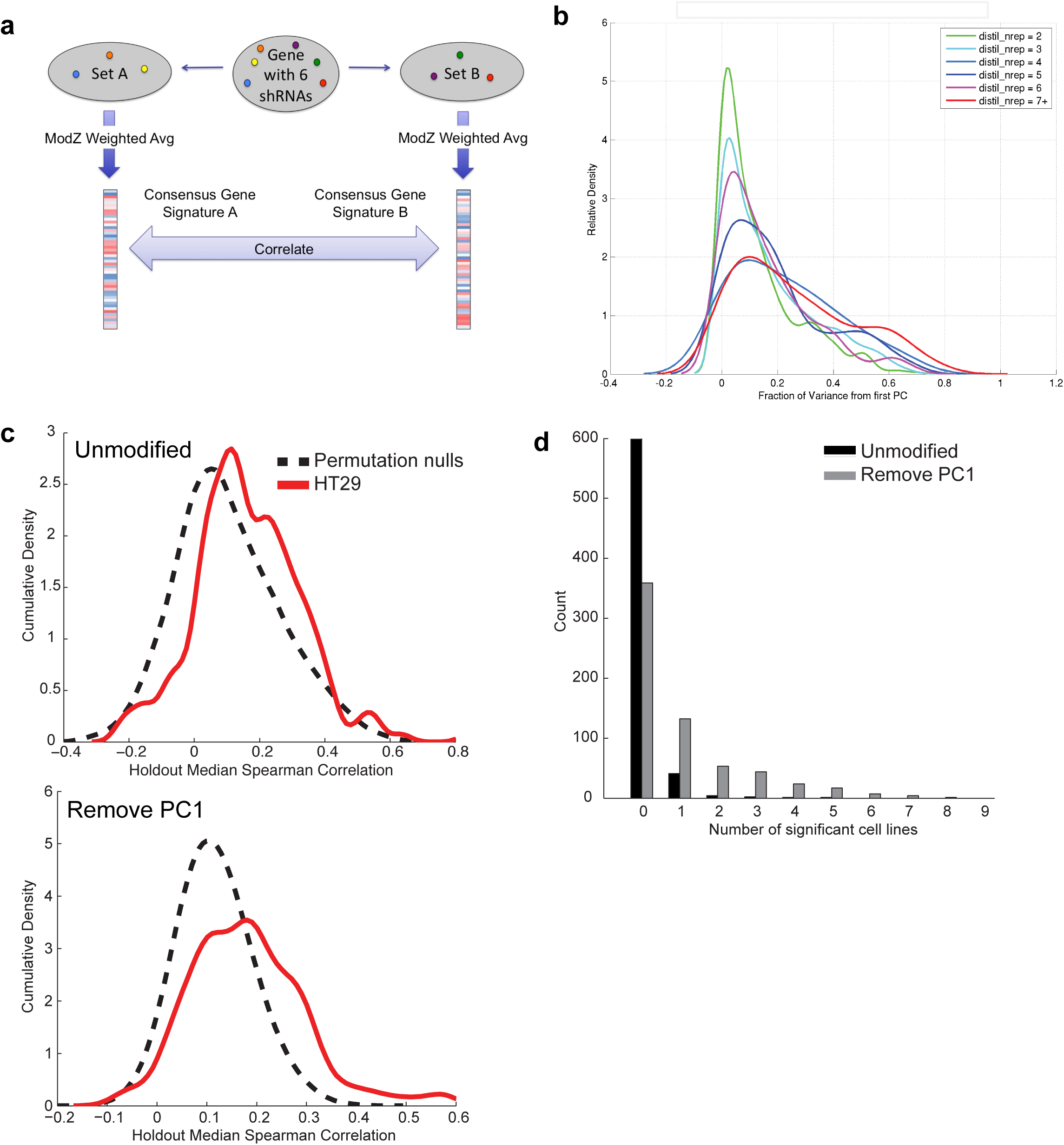
The Consensus Signature and the first Principal Component. a) Schematic of holdout analysis. For genes with 6 or more shRNAs, the Consensus Gene Signature is calculated from subsets of shRNAs, and the resulting CGSs are correlated. This procedure is repeated with different random partitions of shRNAs. b) The increase in correlation variance for random CGSs of increasing numbers of component shRNAs is accompanied by an increase in the relative magnitude of the first principal component, suggesting that it is driving the effect. c) Comparison of holdout correlations to permutations nulls in an example cell line. *Top*: Unmodified signatures; *Bottom*: PC1 removed. Removing PC1 increases the fraction of genes with a statistically significant correlation. d) For each gene, the number of cell lines in which it the CGS passes statistical significance for holdout analysis for unmodified data (black) and for data with the first principal component removed (gray).

**Supplemental Figure 3.**
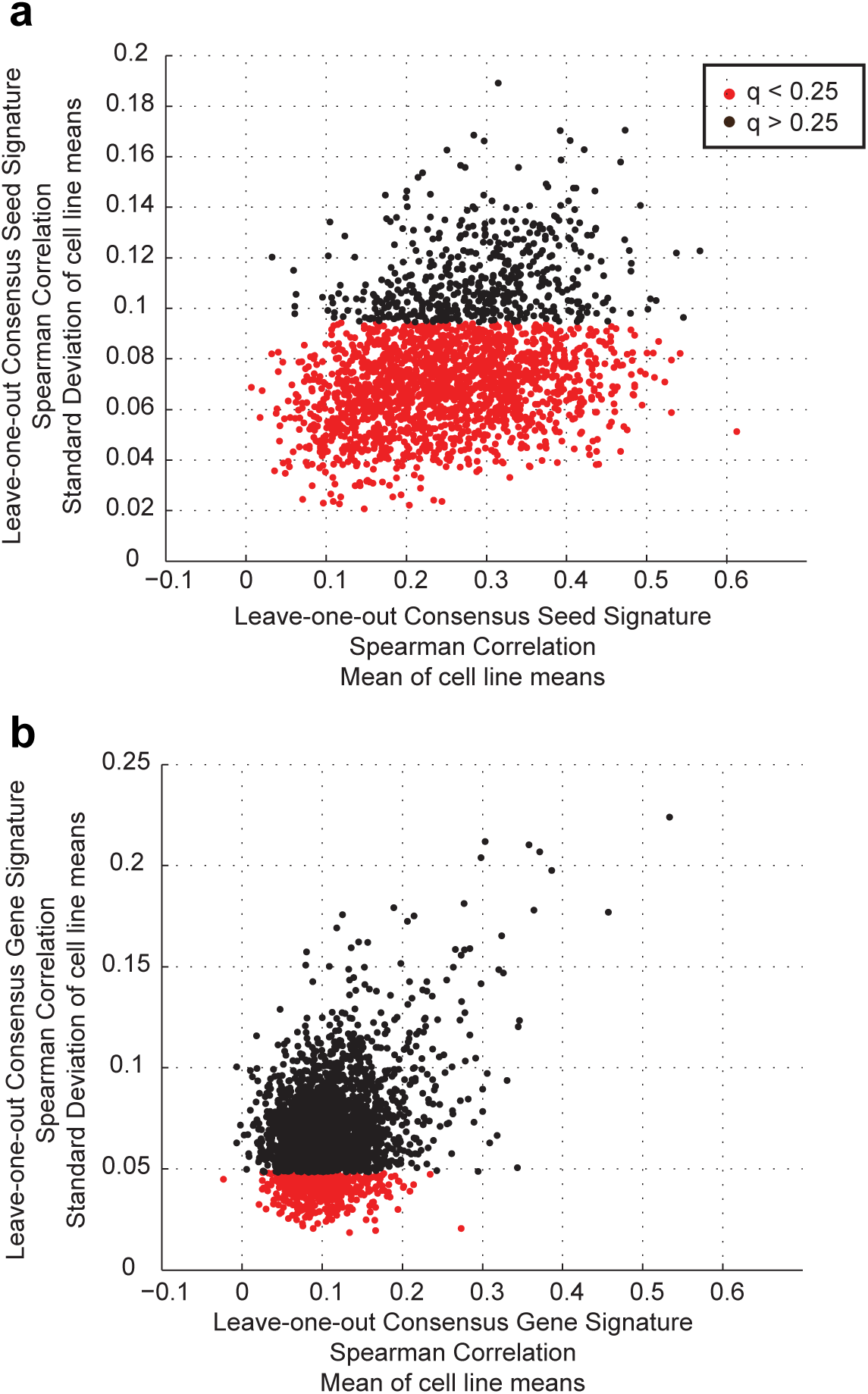
Leave-one-out consensus correlation. For each seed sequence (a) or gene target (b), the mean correlation to the leave-one-out CSS or CGS is calculated within each cell line. The vector of means by cell line for each seed or gene can be compared to the collection of means by cell line for all seeds or genes. Seeds and genes that cause gene expression changes of the same magnitude (not necessarily direction) will tend to have smaller variance than the population, reflecting that consistency. These scatter plots show the mean (x-axis) and standard deviation (y-axis) of the vector of means across the nine cell lines. Those in red are different from the population per the f-test at an FDR of < 25%. Note that the threshold for significance is different for genes and seeds because genes and seeds have different leave-one-one correlation distributions.

**Supplemental Figure 4.**
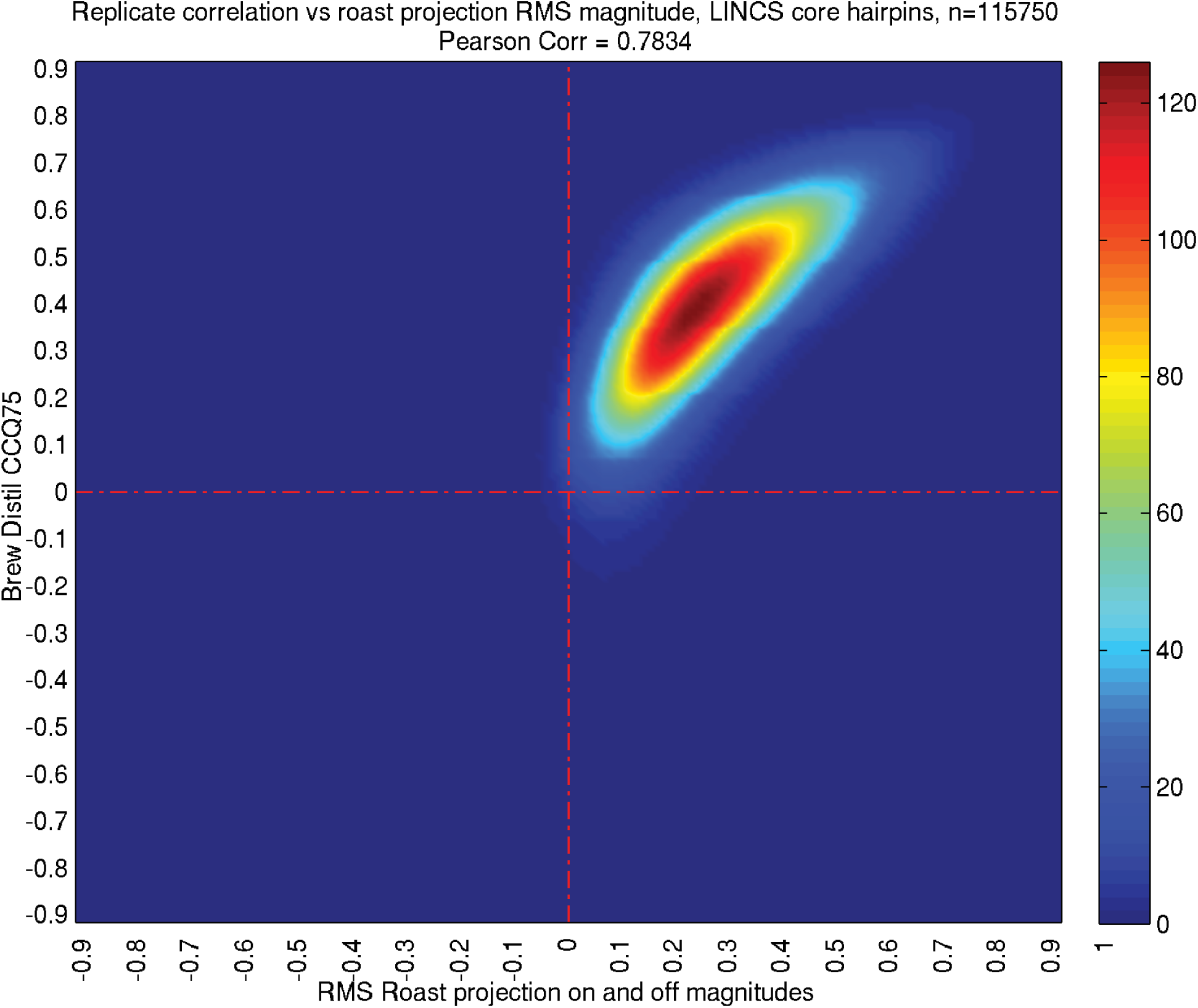
Projection magnitudes recapitulate other measures of reproducibility. The projection algorithm decomposes any particular signature into on-target and residual relative magnitudes. These represent the entirety of the reproducible part of the signature; the remainder is noise. The Pythagorean theorem gives the length of that reproducible part as the square root of the sum of the squares of the two projection magnitudes. This projection length correlates very well with the independently measured replicate correlation. For the set of all shRNAs, the x-axis is the calculated projection length, and the y-axis is the 75^th^ quantile of Spearman correlation among replicates. The two quantities correlate with a Pearson correlation of 0.78, providing a sanity check that the projection outputs are meaningful. The non-linearity is due to the use of the 75^th^ quantile for replicate correlation by convention compared to the use of the mean for projection length.

